# Mapping effective connectivity between the frontal and contralateral primary motor cortex using dual-coil transcranial magnetic stimulation

**DOI:** 10.1101/743351

**Authors:** Karen L. Bunday, James Bonaiuto, Sonia Betti, Guy A. Orban, Marco Davare

**Author notes:** Joint first authors (Karen L. Bunday, James Bonaiuto). Corresponding Author: Karen L Bunday, Psychology, School of Social Sciences, University of Westminster, Room 6.123 (Clipstone) 115 New Cavendish Street London, W1W 6UW.

## Abstract

Cytoarchitectonic, anatomical and electrophysiological studies have divided the frontal cortex into distinct functional subdivisions. Many of these subdivisions are anatomically connected with the contralateral primary motor cortex (M1); however, effective neurophysiological connectivity between these regions is not well defined in humans. Therefore, we aimed to use dual-coil transcranial magnetic stimulation (TMS) to map, with high spatial resolution, the effective connectivity between different frontal regions of the right hemisphere and contralateral M1 (cM1). TMS was applied over the left M1 alone (test pulse) or after a conditioning pulse was applied to different grid points covering the right frontal cortex, while subjects were at rest, prepared an index finger abduction (Prep-ABD) or precision grip (Prep-PG). MEP motor maps were generated by creating synthetic fMRI volumes, including the normalised MEP values at vertices corresponding to the TMS grid locations. These maps were registered to a common atlas, and statistical parametric mapping was used to identify cortical clusters in which stimulation differentially modulated conditioned MEPs across conditions. We found five significant clusters in the frontal cortex. Three clusters in ventral premotor regions (areas 6v and 44) showed significant differential modulations of contralateral MEPs when rest was compared to Prep-ABD and Prep-PG. Two clusters in rostral dorsolateral prefrontal cortex (areas 8Av and 46) showed differential modulation in MEPs when Prep-ABD was compared to Prep-PG. Our findings demonstrate distinct regions that show task-related connectivity whereby interactions between ventral premotor regions and cM1 differentiate between rest and movement preparation and the dorsolateral prefrontal cortex differentiates primarily between preparation of different types of hand movements. These results thus demonstrate the utility of dual-coil TMS and MEP motor maps to define fine-grained sub-regions in the human frontal cortex, which are functional and causally involved in hand movements.

**Highlights:**

- Dual-coil TMS mapping allows defining of fine-grained frontal cortex subdivisions
- Frontal cortex houses multiple areas with different neurophysiological properties
- Interactions between premotor areas and M1 control rest vs movement
- Interactions between prefrontal areas and M1 reflect movement selection

## Introduction

Pioneering studies from neurophysiologist Charles Scott Sherrington provided, using electrical stimulation, the first detailed motor map of the primate motor cortex (Leyton and Sherrington, 1917; reviewed in Lemon, 2008). Interestingly, Sherrington already suggested that the generation of skilled movements ‘certainly involves far wider areas of the cortex than the excitable zone itself’. Using a similar approach in humans, transcranial magnetic stimulation (TMS) studies allowed us to probe the causal role of these ‘silent areas’ in the planning and generation of skilled movements, which are mainly located in the frontal cortex (see Davare et al. 2011 for review). However, due to the vast cortical area to be investigated in humans, the identification of specific frontal cortex subdivisions, which functionally and causally interact with the motor cortex, is still missing. To address this issue, we designed a novel TMS mapping method for investigating the changes in effective connectivity of a large 6×6 cm frontal region with the contralateral motor cortex during skilled hand movements.

The primary motor cortex (M1) and frontal regions of the brain can be distinguished by their cyto-, myelo-, receptor-based architecture and complex array of connections between each other, and other parts of the cortex (Amunts and Zilles, 2015; Fan et al., 2016; Glasser et al., 2016; Luppino and Rizzolatti, 2000; Petrides and Pandya, 1999, 2002). Monkey studies have revealed both anatomical and functional fine-grained subdivisions within the frontal cortex, which have distinctive patterns of connectivity with ipsilateral M1 (Dum and Strick, 1991, 2002; Luppino and Rizzolatti, 2000; Dea et al., 2016; Hamadjida et al. 2016). Although direct callosal projections between contralateral M1 and frontal regions are sparse, if present at all, homotopic and heterotopic callosal projections between other frontal areas are evident in both monkeys and humans (Rouiller et al., 1994; Boussaoud et al., 2005; Lanz et al., 2017; Marconi et al., 2003; de Benediticus et al., 2016). These pathways could be physiologically important and provide information to contralateral M1 (cM1) that helps shape motor commands for voluntary movements, such as grasping and manipulating objects. Modern techniques using magnetic resonance imaging (MRI) in humans support monkey anatomical findings [e.g.(Fan et al., 2016; Glasser et al., 2016)]; however, effective neurophysiological connectivity of these areas is less well defined.

Subdivisions within the frontal cortex are important for the planning and execution of voluntary hand movements, namely the premotor cortex (PM; including area 44), and the dorsolateral prefrontal cortex (DLPFC). Monkey and human ventral and dorsal premotor areas share similar cytoarchitectural features (Kurata, 2018; Petrides and Pandya, 1999, 2002; Petrides et al., 2012). Neuroanatomical tracing techniques in the monkey have revealed that each premotor area has a specific pattern of connections (Dum and Strick, 2002; Kurata and Tanji, 1986). Importantly, premotor regions have homotopic and heterotopic callosal projections to the contralateral hemisphere (Boussaoud et al., 2005; Lanz et al., 2017; Marconi et al., 2003). Single neuron recordings in monkeys have shown that there are anatomically defined subregions of the premotor cortex; the dorsal premotor (PMd) area F2 is particularly involved in the planning and execution of reaching movements and the ventral premotor (PMv) area F5 encodes specific hand actions [for review;(Rizzolatti et al., 2014)]. Human studies also indicate that PMd areas are associated with the preparation of different hand movements (Ariani et al., 2015; Begliomini et al., 2014; Fujiyama et al., 2016; Gallivan et al., 2011), and PMv is associated with object manipulation and grasping (Binkofski et al., 1999; Davare et al., 2008; Ehrsson et al., 2000; Grafton, 2010; Johnson-Frey et al., 2005; Koch et al., 2010).

The DLPFC also has cytoarchitectonically distinct regions that monkeys and humans share (Petrides and Pandya, 1999; Petrides et al., 2012). Although there are no known direct ipsilateral or contralateral connections with M1 (Boussaoud et al., 2005; Fan et al., 2016; Miller, 2000), indirect communication via the premotor cortex and other frontal regions is likely (Boussaoud et al., 2005; Fan et al., 2016; Lanz et al., 2017; Marconi et al., 2003; Miller, 2000). In the monkey, prefrontal cortex neurons code for the internal representations of action goals (Miller, 1999; Wise et al., 1996). Similarly, DLPFC activity is associated with action-related decision making in humans (Bernier et al., 2012; Johnson-Frey et al., 2005; Jueptner et al., 1997; Rowe et al., 2005). Thus, it is possible that anatomical and functional organisation in the monkey and human frontal cortex is similar.

Whereas tract-tracing and microstimulation can be used to probe anatomical and effective connectivity in the monkey brain with high spatial precision, these techniques are not appropriate for non-invasive use in humans. However, effective connectivity between frontal cortical subdivisions and M1 can be probed in the human brain using transcranial magnetic stimulation (TMS) with a conditioning-test paradigm. This involves a conditioning pulse given over the frontal cortex to assess its effect on motor evoked potentials (MEPs) elicited by a test pulse applied over either ipsilateral (Baumer et al., 2009; Civardi et al., 2001; Davare et al., 2008; Koch et al., 2007) or contralateral M1 (Baumer et al., 2006; Buch et al., 2010; Fiori et al., 2017; Koch et al., 2006; Mochizuki et al., 2004). Increases in conditioned MEPs reflect net excitatory influences of the conditioning pulse location on the test pulse site while decreases in conditioned MEPs reveal inhibitory influences. Contralateral frontal cortex - M1 effective connectivity can vary depending on the stimulated site and the performed task (Baumer et al., 2006; Buch et al., 2010; Fiori et al., 2017; Fujiyama et al., 2016; Mochizuki et al., 2004).

The connectivity of subdivisions in the monkey frontal cortex is relatively well known, and while invasive human mapping studies are starting to emerge (Avanzini et al., 2016; Avanzini et al., 2018; Fornia et al., 2018; Vigano et al., 2019), few non-invasive studies in healthy humans have attempted to map large areas of frontal cortex - M1 effective connectivity, or done so during functional tasks (Cattaneo and Barchiesi, 2011; Civardi et al., 2001). We hypothesised that conditioning TMS pulses applied over different subdivisions of the frontal cortex would reveal differential connectivity with cM1 and that varying the condition (rest *vs* preparation of an index finger abduction or a precision grip) would allow us to define the functional properties of clusters of connectivity. To test our hypotheses, we used paired-pulse TMS to examine how conditioning pulses applied to the right frontal cortex affected corticospinal excitability in the opposite M1 during different tasks. Here, we employed a novel approach of mapping physiological motor cortical connectivity using MEPs to create synthetic fMRI volumes, and then leveraging widely used methods from fMRI analysis such as registration to a common template brain and statistical parametric mapping to analyse effective interhemispheric connectivity.

## Methods

### Subjects

Nine right-handed (7 females, 2 males) volunteers aged 22-41 years [age 28±6.3 years (mean±SD)] participated in experiments 1 and 2. Eight subjects participated in both experiments. All subjects were right-handed, with normal, or corrected to normal vision and gave written informed consent. None of the subjects had a history of neurological disease. They were all screened for adverse reaction to TMS using the TMS safety screen questionnaire (Keel et al., 2001). The experimental procedure was approved by the ethics committee of University College London.

### Recordings

Digital conversion and timing of the TMS pulses were performed with a micro CED 1401mk2 unit (Cambridge Electronic Design, Cambridge, UK). Electromyographical (EMG) recordings were made from the first dorsal interosseous (FDI) muscle of the right hand with surface electrodes (belly-tendon montage, Ag-AgCl, 10 mm in diameter). The EMG signal was amplified 1000x, high-pass filtered at 3 Hz, sampled at 5 kHz and stored for off-line analysis (CED 1401 with spike and signal software, Cambridge Electronic Design, Cambridge, UK).

### Transcranial Magnetic Stimulation

Paired-pulse TMS was applied using a Magtim 200 Bistim2 stimulator (Magstim, Whitland, UK) connected to 2 figure-of-eight coils, both produced monophasic waveforms. A standard figure-of-eight coil (9 cm outer diameter, D70) was used for delivering the test pulse (TP) over M1, whereas a custom-made coil (7 cm outer diameter, Magstim custom made) was used for the conditioning pulse (CP) over PM. The test coil was applied tangentially to the scalp with the handle pointing backwards and laterally at a 45° angle to the midline inducing a posterior-anterior (PA) current in the brain (Figure 1A). The coil was systematically moved over the scalp until the optimal “hotspot” was found, identified as the scalp location that reliably produces large motor evoked potentials in the right FDI muscle (Brasil-Neto et al., 1992). The resting motor threshold (RMT), defined as the minimum intensity that induced MEPs of ≥50 μV in 5 out of 10 responses (Rossini et al., 2015; Rothwell et al., 1999), was determined for the right FDI muscle. RMT was 45±9% (mean±SD) of the maximum stimulator output (MSO)]. The intensity of the TP (53±8%) of MSO was set to evoke an MEP of approximately 1 mV peak-to-peak amplitude in the relaxed right FDI muscle. The conditioning pulse was delivered to the right hemisphere with the handle pointing forward with an anterior to posterior (AP) induced current (Figure 1A). The CP was set at 120% (54±12% of MSO) of the RMT (44±9% of MSO) measured with the conditioning TMS coil. The inter-stimulus interval (ISI) between the CP and TP was 8 ms (Baumer et al., 2006; Buch et al., 2010; Neubert et al., 2010; Ni et al., 2009).

**Figure 1.**
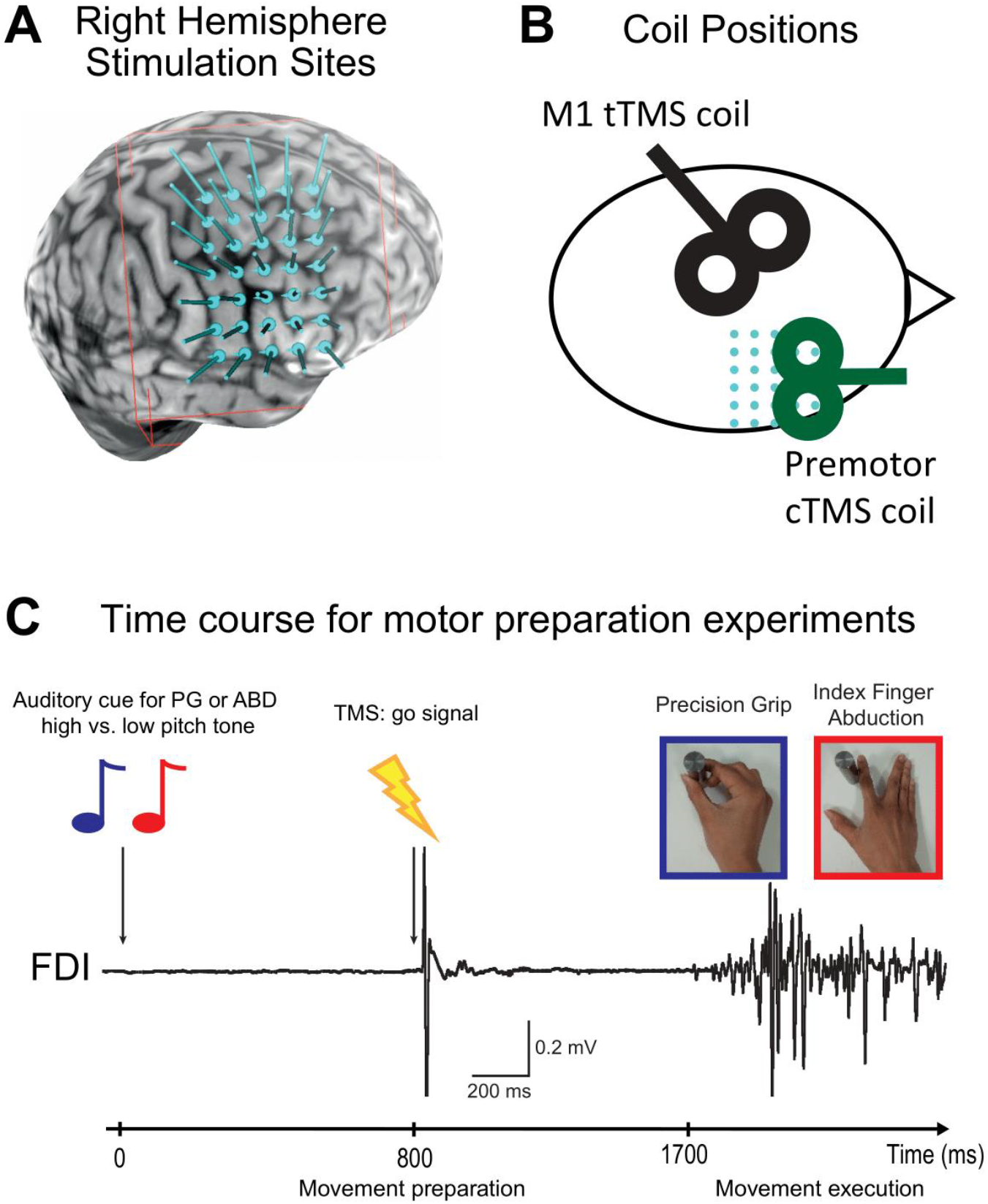
Experimental Setup. **A.** Example of right hemisphere TMS stimulation targets are shown by a 5×7 grid superimposed over the right frontal cortex including the superior frontal sulcus, the central sulcus laterally and the lateral sulcus ventrally (grid point distance: 10 mm). Conditioning TMS coil position was tracked online for precise stimulation over each target. **B.** Coil positions for test and conditioning TMS coils. The test TMS coil was applied tangentially to the scalp over the index finger M1 hotspot with the handle pointing backwards and laterally at a 45° angle to the midline inducing a posterior-anterior (PA) current in the brain. The conditioning TMS coil was applied to the right frontal cortex grid targets with the handle pointing forward with an anterior to posterior (AP) induced current. **C.** Time course for motor preparations experiments. An auditory cue signalled to the subject whether to perform index finger abduction (ABD) or precision grip (PG; high pitch or low pitch, respectively). The TMS TP was triggered 800 ms after the auditory cue and constituted the “go” signal. Subjects subsequently either grasped a cylindrical object between their finger and thumb or abducted the index finger against the object.

The right hemisphere coil locations were determined using an MRI-aligned frameless stereotaxic neuronavigation system (Brainsight, Rogue Research). The Brainsight software 2.2.14 (Rogue Research, Montreal, Canada) was used to reconstruct a three-dimensional image of both the grey matter and the scalp surfaces. A rectangular grid (range: 26-36 targets per grid) was superimposed over the individual’s right hemisphere brain surface using the anatomical MRI. The left border of the grid was aligned to the central sulcus, the top grid border along the superior frontal sulcus and the bottom grid border along the lateral sulcus. The distance between each grid point was 10 mm, and the grid was curved to fit the individual’s brain surface. The grid extended 5-6 grid points rostrally to include the precentral gyrus, middle frontal gyrus, the inferior frontal gyrus including the pars opercularis and pars triangularis. Thus, the grid covered area 4, 6d, 6v, 6a, FEF, 55b, PEF, 6r, i6-8, 8Av, 8C, IFJp, IFJa, 8Ad, 46, p9-46v, IFSp, 44, 45, (Glasser et al., 2016; Figure 1A; Suppl. Fig 1). The conditioning TMS coil was tracked online for precise stimulation of the grid targets; CPs were delivered to each grid point in a randomised order (Brainsight grid point randomisation).

### Experimental Procedure

#### Experiment 1

The first experiment aimed to test the effects of conditioning TMS applied over the right hemisphere on the amplitude of MEPs recorded from the right FDI muscle, elicited by TMS applied over the left motor cortex, at rest (inter-trial interval: 5 s). Participants were comfortably seated in a chair with their hands relaxed on a pillow while TMS pulses were delivered. They were instructed to keep their eyes open and to keep both hands completely relaxed. For each grid point, five conditioned and two unconditioned MEPs were acquired in random order and repeated twice. A total of ten conditioned and four unconditioned MEPs were acquired for each grid point. Therefore, the total number of MEP ranged from 364-504.

#### Experiment 2

The second experiment aimed to test the effects of conditioning TMS applied over the right hemisphere on the amplitude of MEPs recorded from the right FDI muscle, elicited by TMS applied over the left motor cortex, during movement preparation. Here participants were seated comfortably in a chair with their left hand, and arm relaxed, while the right-hand voluntary contracted the FDI by index finger abduction (Prep-ABD) or grasped a solid metal cylindrical object between the index finger and thumb (precision grip; Prep-PG). An auditory cue signalled to the subject whether to perform index finger abduction or precision grip (high pitch or low pitch, respectively). The meaning of the auditory cue (low pitch/high pitch) was counterbalanced between subjects. The TP was triggered 800 ms after the auditory cue and also served as the GO signal (Davare et al., 2009; Prabhu et al., 2007). Thus, TMS was applied during the movement preparation period (Figure 1B). Subjects were instructed to make the movement at their own pace on hearing the TMS click (inter-trial interval: 6.5 s) and to relax in between the trials. At each grid point, five conditioned and two unconditioned MEPs were acquired for preparation of index finger abduction (Prep-ABD) and precision grip (Prep-PG) in a randomised order. Due to the length of the experiment, this procedure was repeated twice in two separate sessions. Thus, over two sessions (1-3 days between sessions), a total of ten conditioned and four unconditioned MEPs were collected for each grid point and each movement condition. Therefore, the total number of MEPs ranged from 364-504 per session.

### Data analysis

The peak-to-peak amplitudes of conditioned and unconditioned MEPs were measured for experiment 1 and 2. Conditioned MEPs were normalised and expressed as a ratio (conditioned MEP/unconditioned MEP). The unconditioned MEP was calculated by a moving average of six unconditioned MEPs (two MEPs from the current grid point + two from the previously acquired grid point + two from the next grid point) rather than to a global average of all unconditioned MEPs from the whole session. For the first and last grid points of a block, the current and next two points and the current and previous two points were used, respectively. This moving averaged unconditioned MEP allowed us to consider small fluctuations in corticospinal excitability throughout the experimental session (2.5-3 hours). MEPs were excluded from analysis if they were preceded by a background EMG activity greater than the resting baseline mean+2SD over a 100 ms window. In addition, for each grid point, trials in which the target error (i.e. the distance between the actual TMS stimulation point and the grid point location on the cortical surface) was 3 SD above the mean (~ > 3 mm) were discarded [Trials excluded per participant (all mean±SD; unless otherwise stated): Expt. 1: 2.95±1.73%; Expt. 2: 2.74±1.91%]. For experiment 2, root mean squared (RMS) EMG activity during the movement was calculated in a fixed window (1000 ms) starting 100 ms after the TMS artefact. In experiment 1, to analyse background EMG activity, a paired t-test was used to compare mean rectified EMG prior to TMS for test and conditioning TMS conditions; for experiment 2, a repeated measure ANOVA (Greenhouse-Geisser correction was used when sphericity could not be assumed) was performed to compare pre and post TMS EMG across sessions and tasks (session 1: Prep-ABD, Prep-PG; session 2: Prep-ABD, Prep-PG). Additionally, a repeated measure ANOVA analysed the variability of test MEP amplitude across experiments and conditions

Analysis of grid based TMS data can be challenging due to heterogeneity in the anatomical structures targeted at each grid location (Fig. 2A). When projected onto a template brain, the row and column gradients of the grid locations in each individual’s native space are still evident (Fig. 2B). Still, there are considerable discrepancies between locations at the group level (Fig. 2C). This spatial heterogeneity in grid point location makes it challenging to perform grid-based group-level analyses. To overcome this difficulty, we developed a novel method of mapping MEP data to individual subject MRI volumes, which could then be mapped to an average template cortical surface for group-level statistics.

**Figure 2.**
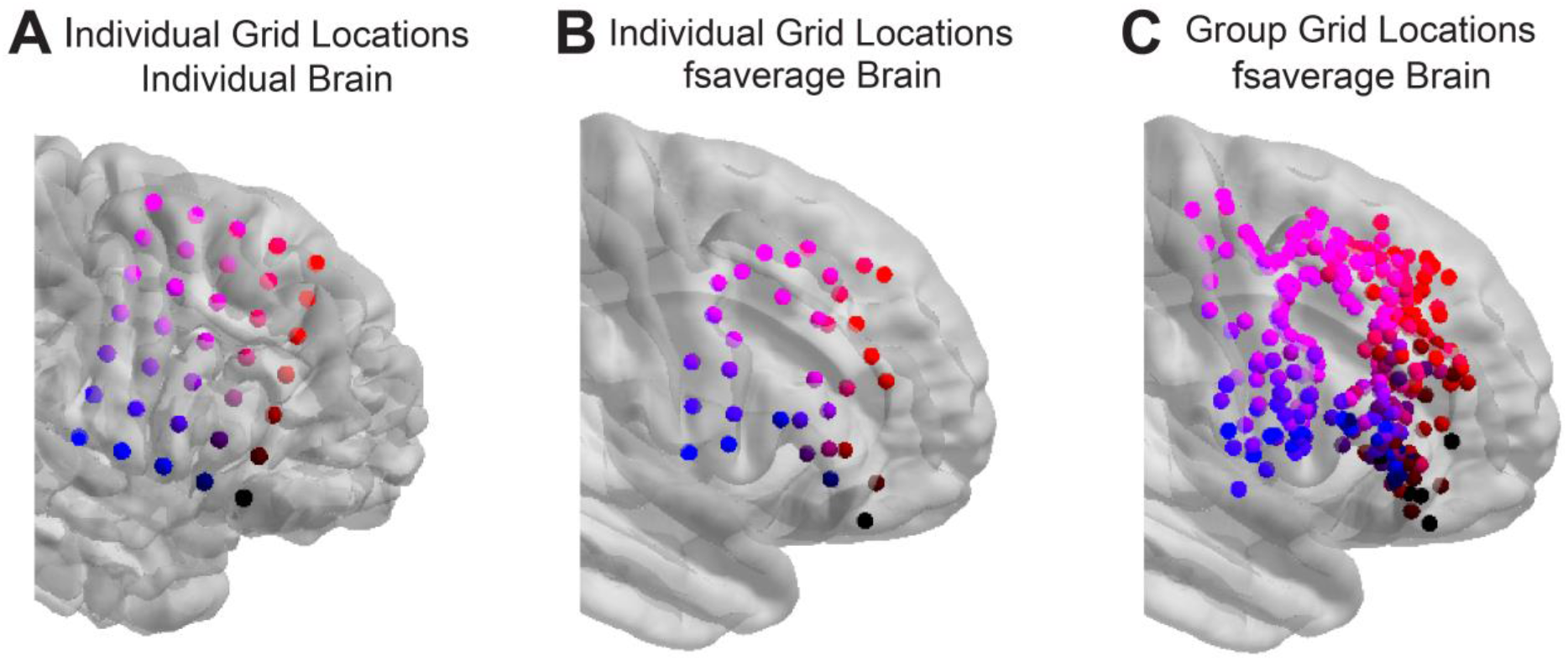
Cortical heterogeneity in individual grid locations. **A.** Each coloured sphere represents the location of a grid point from one participant on their individual brain. Here the colour of each grid point encodes its row and column within the grid. **B.** Grid locations from the same participant, mapped to the FreeSurfer “fsaverage” template brain surface. **C.** There is considerable heterogeneity in grid locations across participants, which is evident then grid locations from all participants are mapped onto the fsaverage brain surface and coloured according to their row and column within each individual subject’s grid.

To achieve this aim, we generated MEP maps for each trial by creating synthetic fMRI volumes from individual MRI brain volumes as follows. First, the normalised MEP values from that trial were attributed to the voxels corresponding to the TMS grid locations (Fig. 3A). These values were then 3D interpolated using SPM12 (Functional Imaging Laboratory, Institute of Neurology, UCL, London, UK) by smoothing them with a Gaussian filter (8mm FWHM) and rescaling the smoothed maps to match the range of normalised MEP values (Fig. 3B). Each participant’s cortical surface was reconstructed from the MRI volumes using FreeSurfer v5.3 (Dale et al., 1999). The interpolated MEP maps (i.e. synthetic fMRI volumes) were then projected onto the individual cortical surfaces (Fig. 3C) and mapped to the FreeSurfer “fsaverage” template brain surface using the inbuilt FreeSurfer function mri_vol2surf (Figure 3D). This mapping was completed via spherical coregistration of the surface topographies (Fischl et al., 1999), which compensates for individual anatomical differences by coregistering individual gyri and sulci.

**Figure 3.**
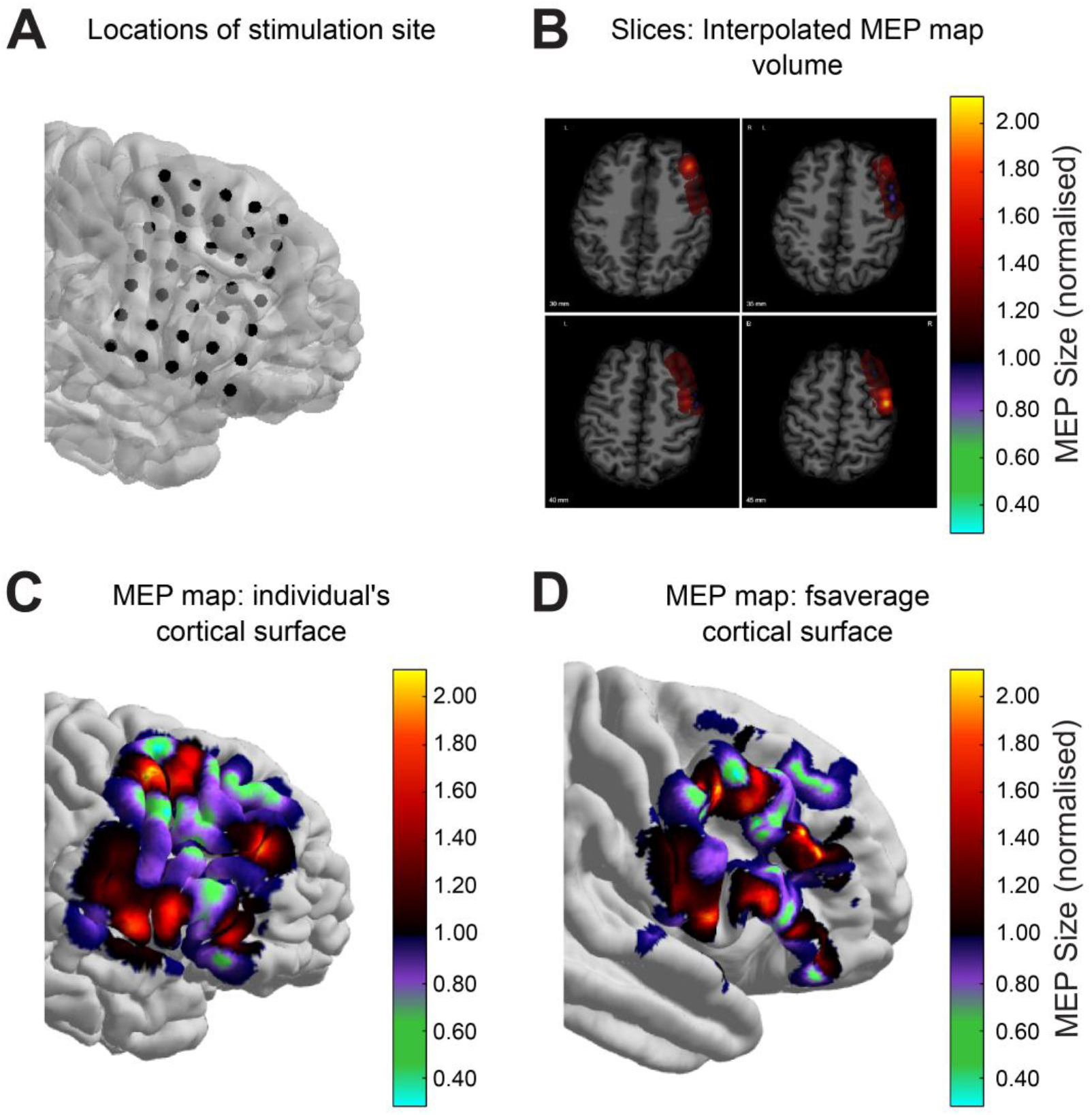
Individual MEP Map Process. **A.** Reconstruction of stimulation location sites for an individual participant. **B.** Axial slices through the interpolated MEP map volume for one trial of the Pre-ABD condition. **C.** The interpolated MEP map for the same trial projected onto the individual’s cortical surface. **D.** The interpolated MEP map surface for this trial aligned to the fsaverage cortical surface.

Once the MEP data were encoded as values in cortical surfaces aligned to a common template brain, we then proceeded to leverage statistical parametric mapping techniques to identify cortical clusters in which stimulation differentially modulated conditioned MEPs across conditions. To identify such clusters of vertices on the cortical template surface, we ran a repeated measures mixed-effects ANOVA using all trials from every subject with a fixed effect of condition (rest, PG, and ABD), and a random effect of subject. We then performed an F-test for the main effect of condition at each vertex within a mask including only vertices with data from all subjects (mask size=7,144 vertices), using an uncorrected p-value threshold of 0.001. This threshold yielded 18 clusters. Next, a cluster size threshold was determined for a p-value threshold of 0.05, corrected for multiple comparisons (family wise error, FWE). This resulted in a cluster threshold of 14 vertices (mean distance between vertices=0.7752mm), yielding 5 significant clusters. The centroid of each cluster was identified, and the cluster labelled by relating its centroid to subdivisions of the cortex based on the Human Connectome Project brain atlas (Glasser et al., 2016), which takes into consideration existing architectonic maps of the frontal cortex (Amiez and Petrides, 2009; Amunts et al., 2010; Amunts et al., 1999; Geyer, 2004; Ongur et al., 2003; Petrides and Pandya, 1999). The anatomical subdivisions and centroids for each cluster are shown in Supplementary Figure S1.

Having identified cortical clusters with differential effective connectivity with contralateral M1, we then ran two analyses within each cluster to compare MEPs between conditions: a surface-based analysis using the MEP map surfaces aligned to the common template brain, and a more classical grid-based analysis. The surface-based analysis used the normalised MEP value averaged over all surface vertices within each cluster as the dependent measure. The grid-based analysis derived the dependent measure by mapping the centroid of each cluster from fsaverage space to each subject’s native space (Fig. 2A, B), and combining the normalised MEP values from the grid points within a threshold distance using a weighted average based on their relative distance to the cluster centroid. The threshold distance was defined as the cluster size significance threshold from the cluster definition (14 vertices) multiplied by the mean distance between vertices (0.7752 mm) to yield a threshold distance of 10.853 mm.

Both the surface- and grid-based analyses used a linear mixed model framework for statistical analysis using R (v3.6.2; R Development Core Team, 2011) and the lme4 package (Bates et al., 2014). For each cluster, a linear mixed model was run with normalised MEP averaged over cluster surface vertice (surface-based analysis) or weighted average of normalised MEPs from grid points within a threshold distance from the cluster centroid (grid-based analysis) as the dependent measure. Note, due to the distance threshold, the number of grid points included the analysis differed by cluster (Table 1). All models included condition as a fixed effect and subject-specific offsets as a random effect. P-values for the fixed effect were obtained using Type III Wald chi-square tests, followed up by planned pairwise comparisons of least square means (Tukey-corrected for multiple comparisons). The normalised MEP values averaged within each cluster (surface-based analysis) and weighted average of normalised MEPs from grid points within a threshold difference from each cluster centroid (grid-based analysis) were compared to 1 (no modulation), using a one-sample t-test.

**Table 1.** Grid points included in grid analysis per subject, per cluster

| Cluster | Mean±SD | Min | Max |
| --- | --- | --- | --- |
| **6v** | 2.89±1.54 | 1 | 4 |
| **44** | 2.33±1.94 | 0 | 5 |
| **44** | 2.89±1.36 | 2 | 4 |
| **8Av** | 2.89±1.27 | 2 | 4 |
| **46** | 2.00±1.41 | 0 | 4 |

Shuffled datasets were generated by permuting the grid locations of each subject, and then generating smoothed MEP maps from the permuted data and aligning them to the fsaverage brain surface. This process was repeated 100 times. We then ran two analyses on the shuffled datasets. The first, aimed at determining how likely we were to get the 5 clusters obtained from the main analysis, ran the cluster definition model on each of the 100 shuffled datasets. The second, aimed at determining the likelihood of getting the condition-dependent results that we obtained for the 5 clusters, averaged the values within the original clusters over the 100 shuffled datasets and ran the follow-up tests from the surface and grid analyses.

## Results

### Control analyses: Test MEPs and background EMG activity

Before considering the main results, it was important to establish standardisation of test MEPs and background muscle activity across the different experiments. We aimed to achieve test MEPs that were similar across both experiments and all conditions; a repeated measures ANOVA revealed no main effect of condition (Expt. 1 - Rest: 1.65±0.63 mV; Expt. 2 – Prep-ABD, Session 1: 1.91±0.55 mV, Session 2: 2.40±1.05 mV; Expt. 2 – Prep-PG Session 1: 1.83±0.68 mV, Session 2: 1.95±0.81 mV; F_(1.9,13.3)_ = 1.61, p=0.24). In experiment 1, background mean rectified EMG prior to TMS was similar across test and conditioning TMS conditions (t_(7)_=1.30, p=0.23). Similarly, in experiment 2, RMS EMG activity before TMS pulses was similar across both sessions (F_(1,7)_ = 0.11, p=0.75) and task (F_(1,7)_ = 0.02, p=0.88). Finally, RMS EMG activity during each movement was similar across sessions (F_(1,7)_ = 0.00, p=0.99) and task (F_(1,7)_ = 4.97, p=0.06). Thus, background EMG both before and after TMS was controlled and therefore did not influence the MEP results.

### MEP analyses

A single TMS pulse was applied to the left motor cortical representation of the FDI muscle either alone or after a conditioning pulse was applied to the right frontal cortex, while subjects were sat at rest or just before an index finger abduction or a precision grip. Normalised MEPs (conditioned MEP/unconditioned MEP) at each grid point over the right frontal cortex were used to create a motor map of physiological interactions between the right frontal cortex and cM1 during each condition (Fig. 3).

At the group level, mean MEP maps appeared to differ markedly between conditions (Figure 4). To find regions where connectivity is significantly different across conditions, we ran repeated measures mixed effects ANOVA at each vertex. Figure 5A shows surface analysis F-value motor maps where significant vertices showed a main effect of condition, thresholded at p<0.001. Thresholding using a cluster size threshold determined for an FWE rate of p<0.05, corrected for multiple comparisons, revealed five significant clusters of vertices (Fig. 5B). The shuffled analysis identified clusters over a wide swath of frontal cortex (Fig. S2A), indicating that these clusters were unlikely to have occurred by chance. This was quantified by computing the exceedance probabilities for a range of percentages of overlap (i.e. the probability that the percentage of vertices shared by two clusters exceeds a given value) between the clusters identified by the shuffled analysis and the 5 clusters identified by the main analysis (Fig. S2B). On average, the probability of finding clusters in the shuffled data which overlapped the original 5 clusters by at least 30% was 0.1.

**Figure 4.**
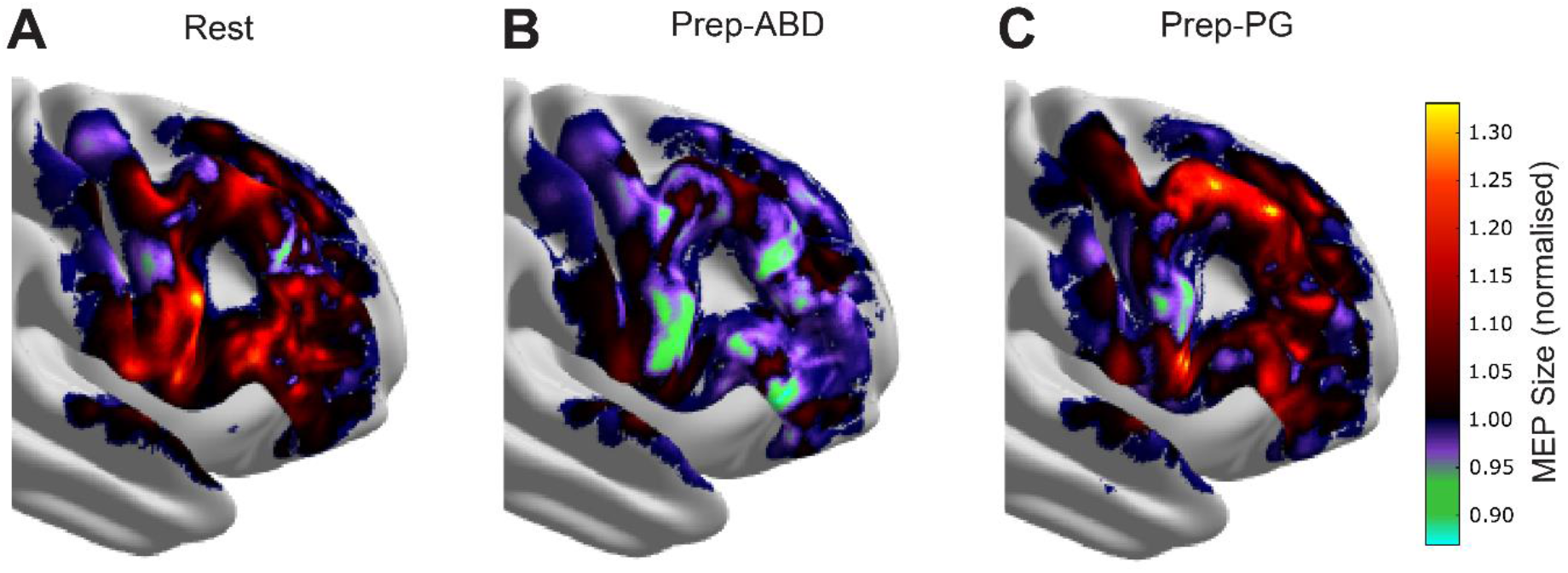
Group MEP Maps. Mean MEP maps on the fsaverage cortical surface for the rest. (**A**), preparation of index finger abduction (Prep-ABD; **B**), and preparation of precision grip (Prep-PG; **C**) conditions. Areas shaded dark red to yellow represents regions where conditioned MEPs were facilitated compared to unconditioned MEPs. Areas shaded purple to blue represent regions where conditioned MEPs were inhibited compared to unconditioned MEPs.

**Figure 5.**
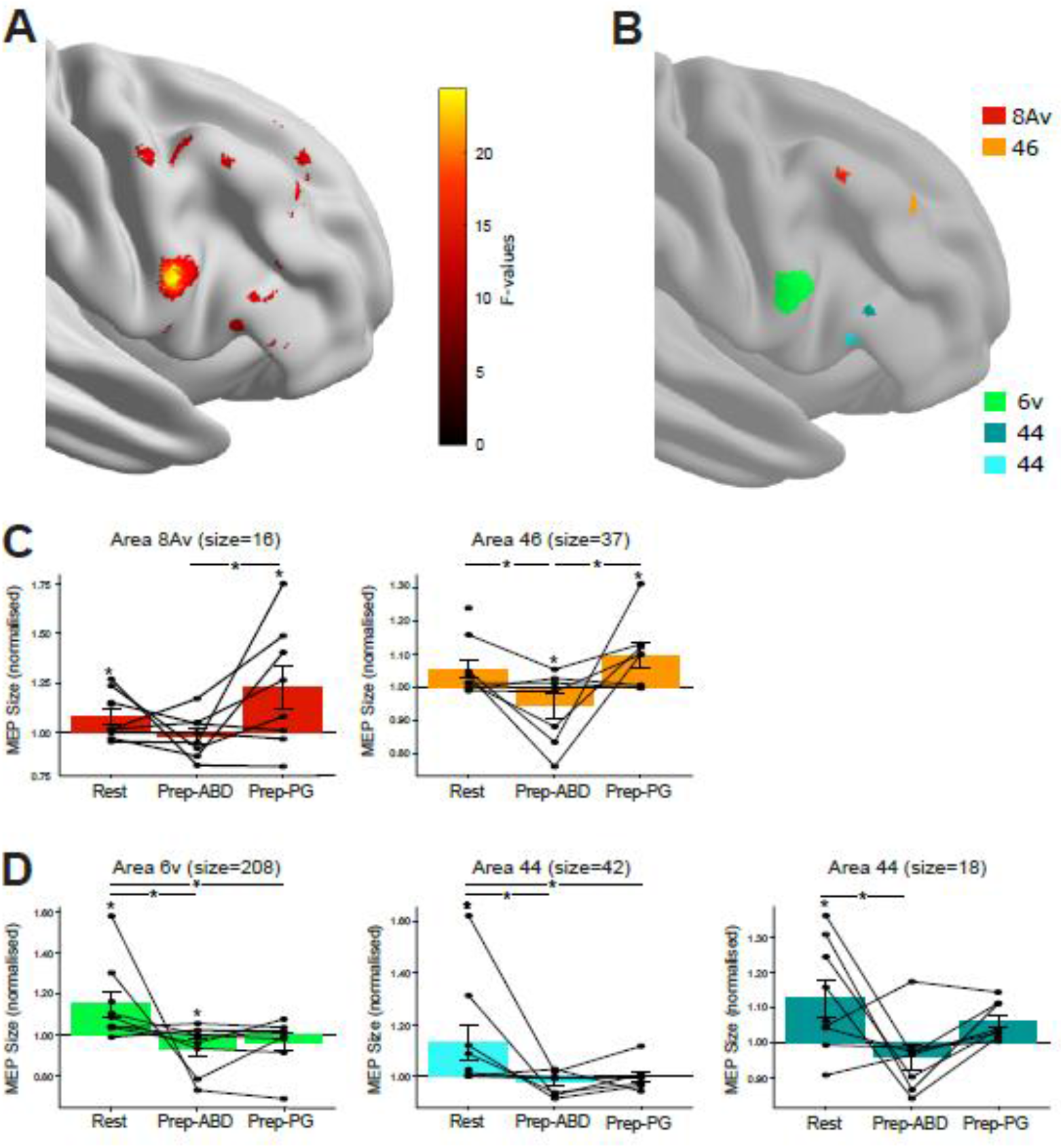
Cluster definition and surface analysis results. **A.** One-way ANOVA F-value map where F-values were thresholded at p<0.05 revealing clusters across the frontal cortex which were differentially modulated by condition. Colour scale represents F-values, where black/dark red represents low F-values, and red/yellow represents high F-values. **B.** Significant clusters found in the frontal cortex after cluster size thresholding. Each cluster colour represents a different region and corresponds to bar graphs in C and D. **C & D.** Bar graphs showing the group normalised MEP size, across all trials and participants, during rest, Prep-ABD and Prep-PG for each cluster. The abscissa shows the condition, and the ordinate shows the normalised conditioned MEP size as a ratio of the unconditioned MEP. Individual participant data points are also shown (filled circles). Each graph corresponds to a cluster shown in B; titles show the area and number of vertices within the cluster. Error bars indicated standard error (SE). *p<0.05.

When comparing the three experimental conditions, these five clusters fell apart in two groups. In three clusters found in ventral premotor regions, testing between conditions, using surface maps and correcting for multiple comparisons, revealed differential effects between rest and Prep-ABD, and between rest and Prep-PG (Fig. 5D). The remaining two clusters in the DLPFC showed significant differences between rest and Prep-ABD and between Prep-ABD and Prep-PG (Fig. 5C). The grid analysis included a mean of 2.84 grid points per subject per cluster (SE = 0.19; see also Table 1; Fig. 6A) and the comparison of MEPs between conditions yielded strikingly similar results to the surface analysis (Fig. 6B, C).

**Figure 6.**
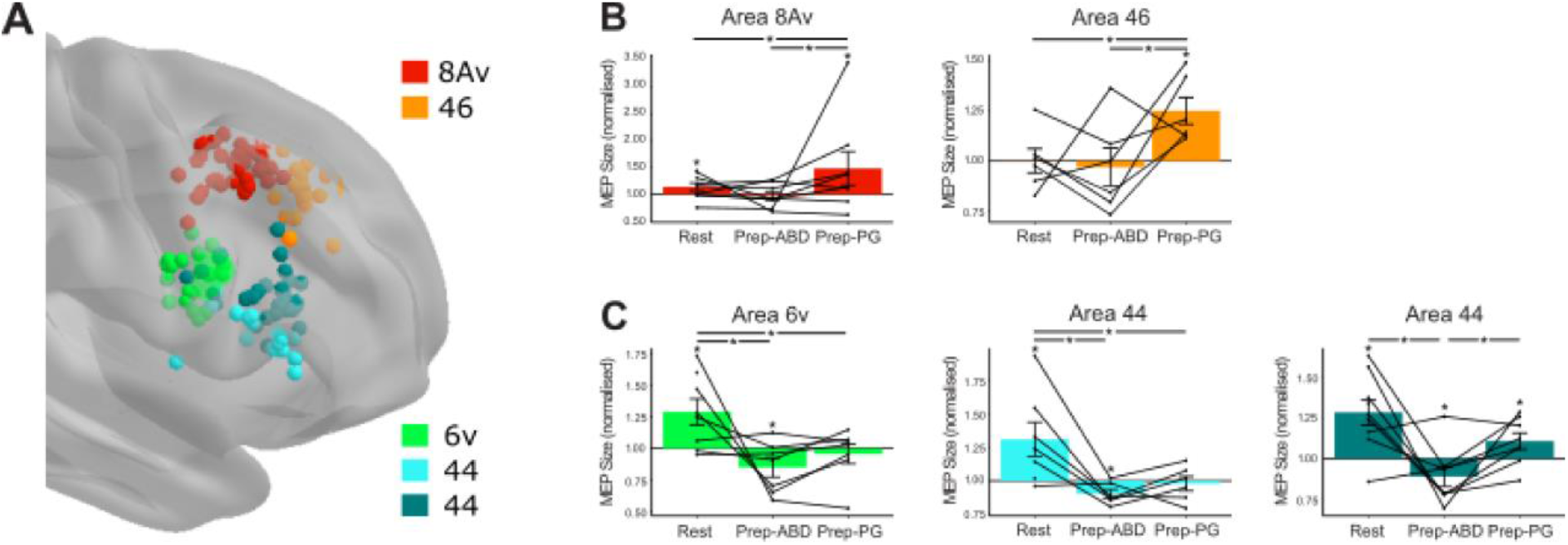
Grid analysis results. **A.** Grid point locations from each subject which were within the distance threshold from a cluster centroid and therefore included in the grid analysis. Each grid location colour represents the cluster to which it was mapped to and corresponds to the bar graphs in B and C. **B & C.** Bar graphs showing the group normalised MEP size, across all trials and participants, during rest, Prep-ABD and Prep-PG for each cluster. The abscissa shows the condition, and the ordinate shows the normalised conditioned MEP size as a ratio of the unconditioned MEP. Individual participant data points included in the analysis are also shown (filled circles). Each graph corresponds to a region shown in A; titles show the cluster area. Error bars indicated standard error (SE). *p<0.05.

### Ventral Premotor Regions of the Frontal Cortex

Overall, significant clusters in ventral premotor regions showed that conditioned MEPs were differentially modulated from rest to movement, but there was no distinction between what movement was to be performed (i.e. index finger abduction or precision grip). All the clusters showed that interactions between premotor areas and cM1 at rest were facilitatory, while only 6v showed significant inhibition when preparing to abduct the index finger. When grid locations were shuffled, no significant main effect of condition was found in these clusters (p>0.05).

#### Area 6v

The largest cluster included 208 significantly active vertices (*X^2^*_(2)_ = 38.09, p<0.001; Table 2; Fig. 5B, D), where the cluster centroid was located in the ventral part of the precentral gyrus, namely area 6 (6v; Fig. S1). This cluster is by far the largest, as it includes nearly two-thirds of the total number (n=321) of active vertices in the five significant clusters (*X^2^*_(5)_ = 130.00, Table 2; p<0.001; Fig. 5B). Within this cluster, the surface-based analysis demonstrated that conditioned MEPs at rest were significantly facilitated compared to Prep-ABD (t_(247)_=−5.63, p<0.001; Fig. 5D) and Prep-PG (t_(247)_=−4.88, p<0.001; Fig. 5D), while index finger abduction and precision grip conditioned MEPs showed similar effects (t_(247)_=−0.74, p=0.742; Fig. 5D). One-sample t-tests showed that conditioned MEPs were significantly facilitated at rest (t_(89)_=4.28, p<0.001; Fig. 5D; Fig. S3) and inhibited during Prep-ABD (t_(79)_=−3.02, p=0.003; Fig. 5D), but not during Prep-PG (t_(79)_=−1.70, p=0.093; Fig. 5D). This pattern of results was also seen in the grid-based analysis (Fig.6C; Table 3).

**Table 2.** One-way ANOVA: Corresponding brain anatomy, brain area (Glasser et al., 2016), active cluster size (total no. of vertices), and MNI coordinates (centroid)

| Neuroanatomy | Area | Cluster Size | X | Y | Z |
| --- | --- | --- | --- | --- | --- |
| Ventral precentral gyrus | 6v | 208 | 61.01 | 4.94 | 28.02 |
| Inferior frontal gyrus - Pars opercularis | 44 | 42 | 56.09 | 19.27 | 12.50 |
| Inferior frontal gyrus - Pars opercularis | 44 | 18 | 54.22 | 23.73 | 21.28 |
| Middle prefrontal gyrus | 8Av | 16 | 43.48 | 14.01 | 51.14 |
| Middle prefrontal gyrus | 46 | 37 | 34.26 | 31.46 | 40.72 |

**Table 3.**
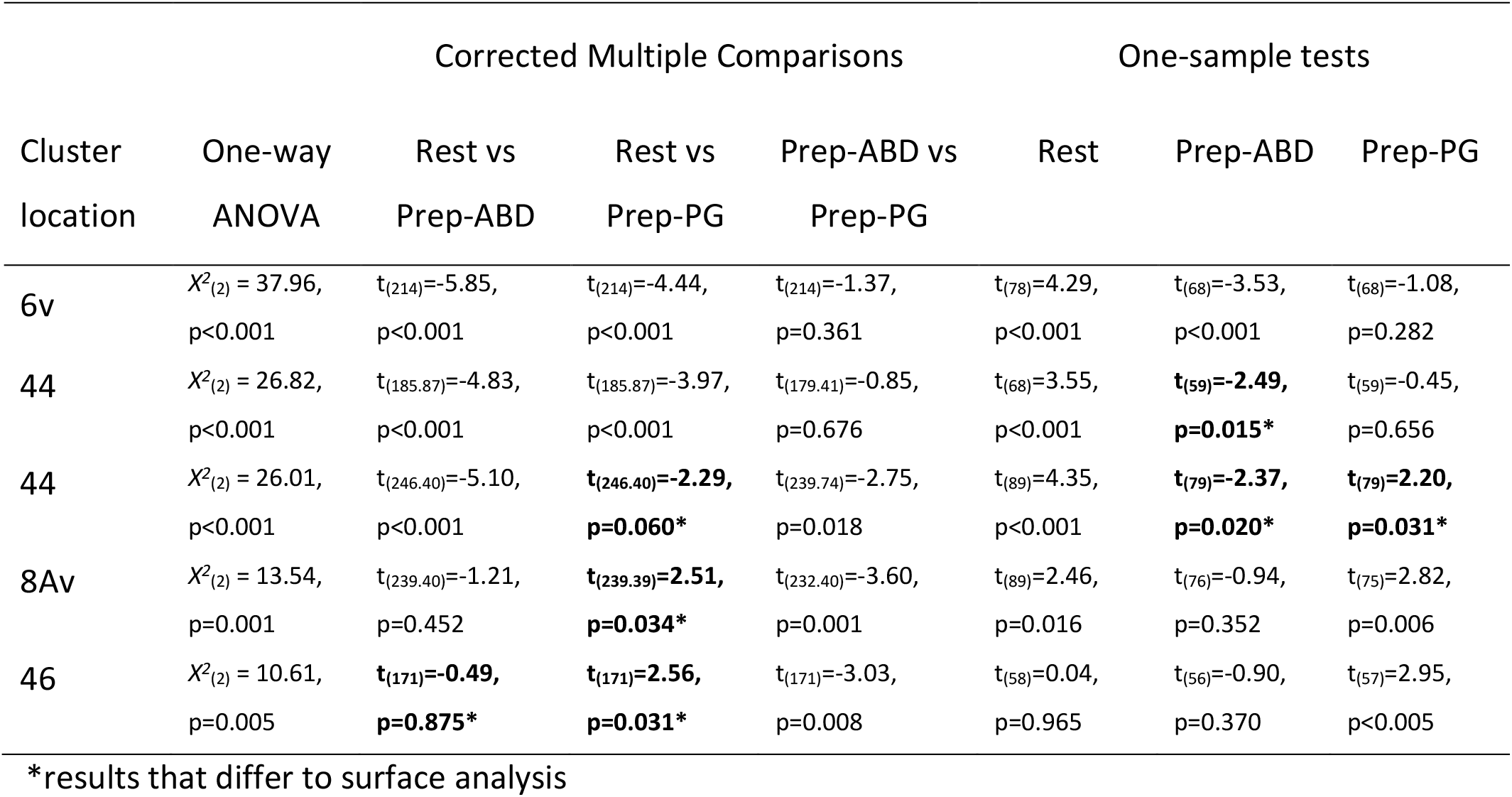
Grid analysis one-way ANOVA and one-sample t-test results

#### Area 44

Two significantly active clusters (Fig. 5B, D; Table 2) were found in the inferior frontal gyrus (IFG), specifically the pars opercularis. The centroids of both clusters were found within area 44 (Fig. S1). Similar to area 6v, the larger, more ventral, significant cluster (42 vertices; *X^2^*_(2)_ = 19.03, p<0.001; Fig. 5B, D; Table 2) revealed that conditioned MEPs at rest were significantly facilitated compared to Prep-ABD (t_(245.91)_=−4.02, p<0.001; Fig. 5D) and Prep-PG (t_(245.91)_=−3.43, p=0.002; Fig. 5D), but no difference was found between Prep-ABD and Prep-PG (t_(239.13)_=−0.58, p=0.830; Fig. 5D). One-sample t-tests showed that contralateral conditioned MEPs were significantly facilitated at rest (t_(89)_=3.17, p=0.002; Fig. 5D), but showed no change during Prep-ABD (t_(79)_=−1.42, p=0.159; Fig. 5D) or Prep-PG (t_(79)_=0.02, p=0.985; Fig. 5D). Similar results were seen in the grid-based analysis. However, Prep-ABD MEPs were significantly inhibited (Fig. 6C; Table 3).

The smaller, slight more rostral significant cluster (18 vertices; *X^2^*_(2)_ = 10.57, p=0.005; Fig. 5B, D; Table 2; Fig. S1), showed that conditioned MEPs at rest were significantly facilitated compared to Prep-ABD (t_(246.22)_=−3.24, p=0.004; Fig. 5D), but other comparisons were not significant (p>0.05; Fig. 5D). One-sample t-tests revealed that contralateral conditioned MEPs were significantly facilitated at rest (t_(89)_=2.83, p=0.006; Fig. 5D), but showed no change during Prep-ABD (t_(79)_=−1.36, p=0.179; Fig. 5D) and a trend towards facilitation during Prep-PG (t_(79)_=1.92, p=0.059; Fig. 5D). The grid-based analysis showed some similar effects, but conditioned MEPs were also differentially modulated during Prep-ABD, and Prep-PG (Fig. 6C; Table 3), where Prep-ABD and Prep-PG conditioned MEPs were significantly inhibited and facilitated (Fig. 6C; Table 3), respectively.

### Dorsolateral Prefrontal Cortex

Significant clusters found in the DLPFC showed a different pattern of effects compared to the premotor regions. Here, conditioned MEPs in both clusters were differentially modulated when preparing for the two different types of movement and significantly facilitated during Prep-PG. The more rostral of the two clusters (area 46) showed further differential modulation between rest and Prep-ABD and significant inhibition during Prep-ABD. When grid locations were shuffled, no significant main effect of condition was found in these clusters (p>0.05).

#### Area 8Av

A significant cluster was found on the crown of the middle frontal gyrus and consisted of 16 significantly active vertices (*X^2^*_(2)_ = 15.27, p<0.001; Fig. 5B, C; Table 2). Its cluster centroid was found in the area 8Av (Fig. S1). Within this cluster, conditioned MEPs were significantly facilitated during Prep-PG compared to Prep-ABD (t_(239.71)_=−3.89, p<0.001; Fig. 5C) and a trend towards significance when compared to rest (t_(246.39)_=2.32, p=0.055; Fig. 5C). No significant difference was found between rest and Prep-ABD (t_(246.39)_=−1.66, p=0.220; Fig. 5C). One-sample tests revealed that contralateral conditioned MEPs were significantly facilitated at rest (t_(89)_ = 2.21, p=0.030; Fig. 5C) and during Prep-PG (t_(79)_ = 3.58, p<0.001; Fig. 5C), but no change in MEPs during Prep-ABD (t_(79)_ = −0.84, p=0.406; Fig. 5C). This pattern of results was also seen in the grid-based analysis, however, the difference between rest and Prep-PG MEPs was significant (Fig.6B; Table 3).

#### Area 46

A larger significant cluster of 37 vertices was found in a more rostral region of the middle frontal gyrus (*X^2^*_(2)_ = 12.81, p=0.002; Fig. 5B, C; Table 2), with its centroid located in area 46 (Fig. S1). Like area 8Av, conditioned MEPs at during Prep-PG were facilitated compared to Prep-ABD (t_(236.93)_=−3.452, p=0.002; Fig. 5C). Additionally, rest conditioned MEPs were significantly facilitated compared to Prep-ABD (t_(245.80)_=−2.57, p=0.028; Fig. 5C). No significant difference was found between rest and Prep-PG (t_(245.80)_=0.97, p=0.599; Fig. 5C). One-sample tests revealed that contralateral conditioned MEPs were significantly inhibited during Prep-ABD (t_(79)_ = −2.82, p=0.006; Fig. 5C), but significantly facilitated during Prep-PG (t_(79)_ = 2.62, p=0.010; Fig. 5C), but there was no change in conditioned MEP size at rest (t_(89)_ = 1.73, p=0.087; Fig. 5C). The grid-based analysis showed some similar results; however, there was no significant differential effect between rest and Prep-ABD conditioned MEPs (Fig. 6B; Table 3). Instead Prep-PG conditioned MEPs were significantly facilitated compared to rest (Fig. 6B; Table 3). Additionally, Prep-ABD conditioned MEPs were not significantly inhibited (Fig. 6B; Table 3).

## Discussion

Our study used dual-coil TMS to examine interactions between the right frontal cortex and cM1. Based on their different effective connectivity profiles, we identified five frontal subdivisions that showed task-dependent differential modulation of corticospinal excitability. Specifically, three clusters found in the ventral premotor regions of the frontal cortex (areas 6v and 44) showed a net facilitatory influence on cM1 MEPs at rest compared to preparation of index finger abduction and precision grip. In area 6v, significant facilitation switched to inhibition during preparation of index finger abduction. Two clusters found in rostral DLPFC, namely areas 8Av and 46, showed a net facilitatory influence on cM1 MEPs during precisions grip compared to index finger abduction. Area 46 showed a significant switch from facilitation during preparation of precision grip to inhibition during index finger abduction. Ultimately, our results show distinct regions of task-related interactions wherein ventral premotor regions interact with M1 by differentiating between rest and preparing to move, but DLPFC regions differentiate between preparing different types of hand movement. Overall, these results demonstrate, for the first time non-invasively in humans, the nature of neurophysiological interactions (i.e. inhibition *vs* facilitation) between multiple frontal subdivisions and cM1 and how these interactions change depending on the movement context.

### Ventral Premotor Cortex

Changes revealed in the net motor output provide insights into how different populations of neurons respond, and how different cortical pathways are recruited, in different contexts. Our results showed that there were three clusters of vertices within ventral PM regions (6v and 44) which differentially modulate net cM1 output when comparing rest to index finger abduction and precision grip preparation. All three regions showed a significant facilitatory influence on the cM1 at rest. However, the switch towards inhibition during index finger abduction preparation was only found in area 6v. A limited number of TMS studies have examined PMv connectivity with cM1 at rest, although some evidence suggests that the influence of PMv on cM1 can be inhibitory (Fiori et al., 2017). However, the longer ISIs used in Fiori et al. (2017) study (e.g. 40, 150 ms) are likely to involve other cortical and subcortical networks. During movement preparation, facilitatory effects of PMv on cM1 have been demonstrated (Buch et al., 2010), although the direction of these effects can be both timing and task-dependent (Buch et al., 2010; Neubert et al., 2010). Despite a similar region to our 6v being stimulated, our results stand in contrast to those of Buch et al. (2010) which revealed a facilitatory influence of PMv on cM1 during movement preparation. However, methodological differences could explain this discrepancy since TMS was delivered much earlier after the movement cue in the Buch et al. (2010) study and they also used a lower stimulation intensity (110% of RMT). Nonetheless, our and Buch et al. (2010) results might suggest that the influence of PMv on the cM1 changes during the preparation period before movement execution, is intensity-dependent, and depends on the type of movement to be made.

PMv forms part of the dorsolateral grasping circuit (Grafton, 2010; Grol et al., 2007). Imaging studies show that PMv regions similar to our study are involved in object manipulation (Binkofski et al., 1999; Ehrsson et al., 2000; Johnson-Frey et al., 2005), execution and planning of precision grasping (Grol et al., 2007). However, we show effective connectivity from the right hemisphere, and the latter studies only show activation in the left hemisphere. While the organisation of right and left area 6 (our largest cluster) are similar (Avanzini et al., 2018), the right and left IFG are different, where right IFG has in fact been associated with response inhibition during motor execution (Aron et al., 2014). Analysis of whole-brain activation patterns across a large database of neuroimaging experiments has shown that area 6 related to action imagination and action execution, whereas area 44 was associated with memory, encoding and spatial attention (Hartwigsen et al., 2019), suggesting different functions in these regions. In the monkey, the ventral premotor cortex likely consists of areas F4 and F5 (Rizzolatti et al., 2014; Kurata, 2018) and the human PMv corresponds at least partially to F5. Neurons in these areas code for a variety of grasping actions (Fluet et al. 2010; Rizzolatti et al., 2014; Bruni et al., 2017), but also the goal and temporal aspects of the action (Jeannerod et al., 1995). Despite this detailed functional evidence, our study investigated three simple conditions; similar effects for rest and the preparation of the two movement tasks were seen in both area 6v and 44, demonstrating that TMS given at 800 ms after the auditory cue was able to reveal differential encoding of movement preparation compared to rest, but this encoding did not discriminate between the types of hand action. Nonetheless, it is interesting that the largest cluster of active vertices was found in area 6v. Given that this region is heavily involved in the control of hand movements (Fluet et al. 2010; Kurata, 2018), a large area of differential net connectivity here may not be surprising.

Our results must be considered in relation to potential anatomical pathways. There are known ipsilateral projections from PM to M1 in the monkey (Dum and Strick, 2002; Rizzolatti et al., 2014), which also may exist in the human (Davare et al., 2008; Davare et al., 2009), so it possible that information flows first to iM1 and then to cM1 (i.e. iPM – iM1 – cM1). Alternatively, PM transcallosal projections could also transfer this information to the contralateral frontal regions. For instance, studies show that there are dense homotopic and heterotopic callosal connections between the right and left dorsal and ventral PM, with sparser heterotopic connections from other motor and non-motor regions (Boussaoud et al., 2005; Lanz et al., 2017; de Benedictus et al., 2016). In support, functional connectivity data shows links between right area 6 and a network that includes the left area 44 and right area 44 connects to a network that includes left area 44 and 6 (Hartwigsen et al., 2019). This provides the current study with two main possible pathways for the net cM1 output, namely iPM – cM1 or iPM – cPM – cM1. Since callosal projections to cM1 from premotor regions are sparse or not present in the monkey or human (Rouiller et al., 1994), and are likely to be faster than our ISI might allow, we suggest that the net cM1 output in our study likely results from information being passed via the contralateral PM (i.e. iPM – cPM – cM1). Our results favour this pathway since M1 transcallosal pathways project onto local circuitry before finally connecting with cM1 corticospinal neurons at ISIs of 8 and 10 ms (Di Lazzaro et al., 1999; Lee et al., 2007; Ni et al., 2009), which is comparable to our ISI.

### Dorsolateral Prefrontal Cortex

We found two significant clusters in regions of the DLPFC, namely areas 8Av and 46. These clusters revealed somewhat similar results in that stimulation of these regions differentially modulated cM1 net output during index finger abduction and precision grip preparation. Area 8Av, but not 46, had a facilitatory effect on cM1 at rest. Similar TMS studies have shown no effect of DLPFC stimulation on cM1 at rest (Mochizuki et al., 2004; Ni et al., 2009). The location in these studies is closer to our area 46, suggesting a gradient of net facilitation where more posterior regions show greater facilitation at rest. A significant inhibitory effect was only found in area 46 during index finger abduction preparation, while both 8Av and 46 switched back to facilitation during precision grip preparation. These results perhaps illuminate the role of DLPFC choosing the correct movement to make during the preparation phase. A facilitatory influence of DLPFC on cM1 has been demonstrated when subjects prepare to make bilateral movements (Fujiyama et al., 2016). Moreover, repetitive TMS over the lateral prefrontal cortex reduces inhibition in the opposite M1 of a non-selected effector during the planning of finger movements in a choice reaction task (Duque et al., 2012). The authors suggest that this region is important for selecting the appropriate response (Duque et al., 2012). Similarly, imaging studies have shown that the DLPFC is important in action-based decision making (Bernier et al., 2012; Johnson-Frey et al., 2005; Jueptner et al., 1997; Rowe et al., 2008). Furthermore, neural activity in this area is related to behavioural goals (Yamagata et al., 2012) and the integration of information for action planning (Hoshi and Tanji, 2004; Tanji and Hoshi, 2008).

Anatomical studies have shown that there are no known ipsilateral connections between DLPFC and M1 (Miller and Cohen, 2001). However, this does not negate the fact that stimulation of this region can affect net M1 output (Hasan et al., 2013), possibly via PMd or other frontal regions (Lu et al., 1994; Luppino et al., 1993; Miller, 2000; Bakit et al.; 2020; Schulz et al.; 2019). A recent study showed that caudal and middle regions of area 46 ipsilaterally project to both dorsal and ventral premotor regions, as well as area 8 (Borra et al., 2019). Similarly, in the human, there are known intralobular connections between DLPFC and PMd and PMv (inferior frontal gyrus – pars triangularis; (Catani et al., 2012) that information could be passed between before being relayed to ipsilateral M1 or cM1 (i.e. DLPFC – iPMv/iPMd – iM1 – cM1 or DLPFC – iPMv/iPMd – cPMv/cPMd – cM1). As this route is quite long and connects four separate regions, it is unlikely to be compatible with the ISI used in our study. Alternatively, information concerning action choice could be relayed from DLPFC to cPMd then to cM1, since some monkey studies show strong projections from the prefrontal cortex to cPMd (Boussaoud et al., 2005). Although other studies show that DLPFC and PM projections are somewhat sparser, the number projections are non-negligible and thus still probably functionally relevant (Lanz et al., 2017; Marconi et al., 2003). In support, diffusion tensor imaging in humans suggests there is significant connectivity between the middle frontal and the precentral gyrus (Jarbo et al., 2012). Whichever the route used to convey information from DLPFC to cM1, stimulation of the area affects net contralateral motor output in a task-dependent manner.

Our results show two distinct patterns of net frontal cortex connectivity with cM1. While four of the five clusters show significant net facilitation at rest, the clusters diverge during the preparation of index finger abduction and precision grip. Premotor regions distinguish between rest and movement preparation in general, while DLPFC shows significant differential modulation during the different movements. These results suggest premotor region clusters show differential net connectivity which code for whether to move irrespective of the type of hand movement and DLPFC clusters differentiate between which movement should be made. This pattern of net connectivity is consistent with the affordance competition hypothesis, whereby neural populations in the frontal regions representing potential movements compete between each other (Cisek, 2007). In this context, premotor regions might prepare plans for both movements DLPFC then selects the action in cM1.

### Methodological considerations

In this study, we have presented a novel method of non-invasive evaluation of effective connectivity within the human motor system. This involves the creation of synthetic fMRI volumes of MEP modulation, which are then mapped to a template cortical surface in order to leverage statistical parametric mapping techniques for group-level analyses. We show that grid-based analysis within this space supports the results of the surface-based analysis, justifying analyses of MEP maps on cortical surfaces mapped to a common template surface in future studies. This approach allows for high-density stimulation-based studies of effective connectivity while accounting for anatomical heterogeneity between subjects. However, this study has several limitations that warrant discussions such as the sample size, use of a single ISI, and the anatomical target of TMS.

Due to the nature and length of mapping based TMS studies, our sample size was small. Nonetheless, our study used many stimulation sites in conjunction with a mixed model approach which meant we were able to use every MEP rather than subject-averaged data. Thus, despite the sample size limitation, these combined methods allowed us to explore frontal cortex to contralateral net connectivity non-invasively.

Evidence reveals that there are several potential pathways in which stimulation of premotor and DLPFC regions can affect contralateral motor output. Most of these pathways involve a number of synaptic connections before arriving at corticospinal neurons. Thus an ISI of 8 ms could seem relatively short; however, it has been suggested that in other established interhemispheric interactions transcallosal projections are likely to synapse onto local circuitry before information arrives at corticospinal neurons in layer V (Di Lazzaro et al., 1999; Ferbert et al., 1992; Lee et al., 2007; Ni et al., 2009). In those subjects where small effects are seen it is possible that a longer ISI could have improved effective connectivity; however, it was not possible to test several ISIs, given our experiment already contained a large number of conditions.

Modelling research has shown that TMS predominately affects the gyral crown underneath the focal point of the coil (i.e. the centre of a figure of eight coil), but surrounding areas can also be affected depending on the gyral architecture (Opitz et al., 2013; Thielscher et al., 2011). Thus, it could be possible that neighbouring clusters are artefacts of nearby TMS depending on the surrounding gyral structure. However, since we used a thresholded cluster size and most of our significant clusters were centimetres away from each other, this is less likely.

## Conclusions

We present a novel method for high-density analysis of effective connectivity using MEP maps registered to the cortical surface of a template brain. Using this method, we show that it is possible to use non-invasive dual-coil TMS to define frontal subdivisions that are important for the control of voluntary movement and shed light on the physiological nature of their interactions with M1. In line with previous research, our results reveal that premotor regions code for whether to move regardless of movement type and the DLPFC codes for which movement should be made. These effects show distinct areas, where the former differential effects (i.e. whether to move or not) are located in ventral frontal regions, and dorsolateral regions encode for different movement goals.

## Acknowledgements

This work was supported by the Biotechnology and Biological Sciences Research Council (BBSRC; Grant numbers: BB/N001370/1 and BB/J014184/1 to M. Davare), the European Union’s Horizon 2020 research and innovation programme (ERC consolidator grant agreement number 864550 to J. Bonaiuto) and the European Research Council (ERC - Parietal action; Grant number 323606 to GA. Orban).

## Supplementary Figures

**Figure S1.**
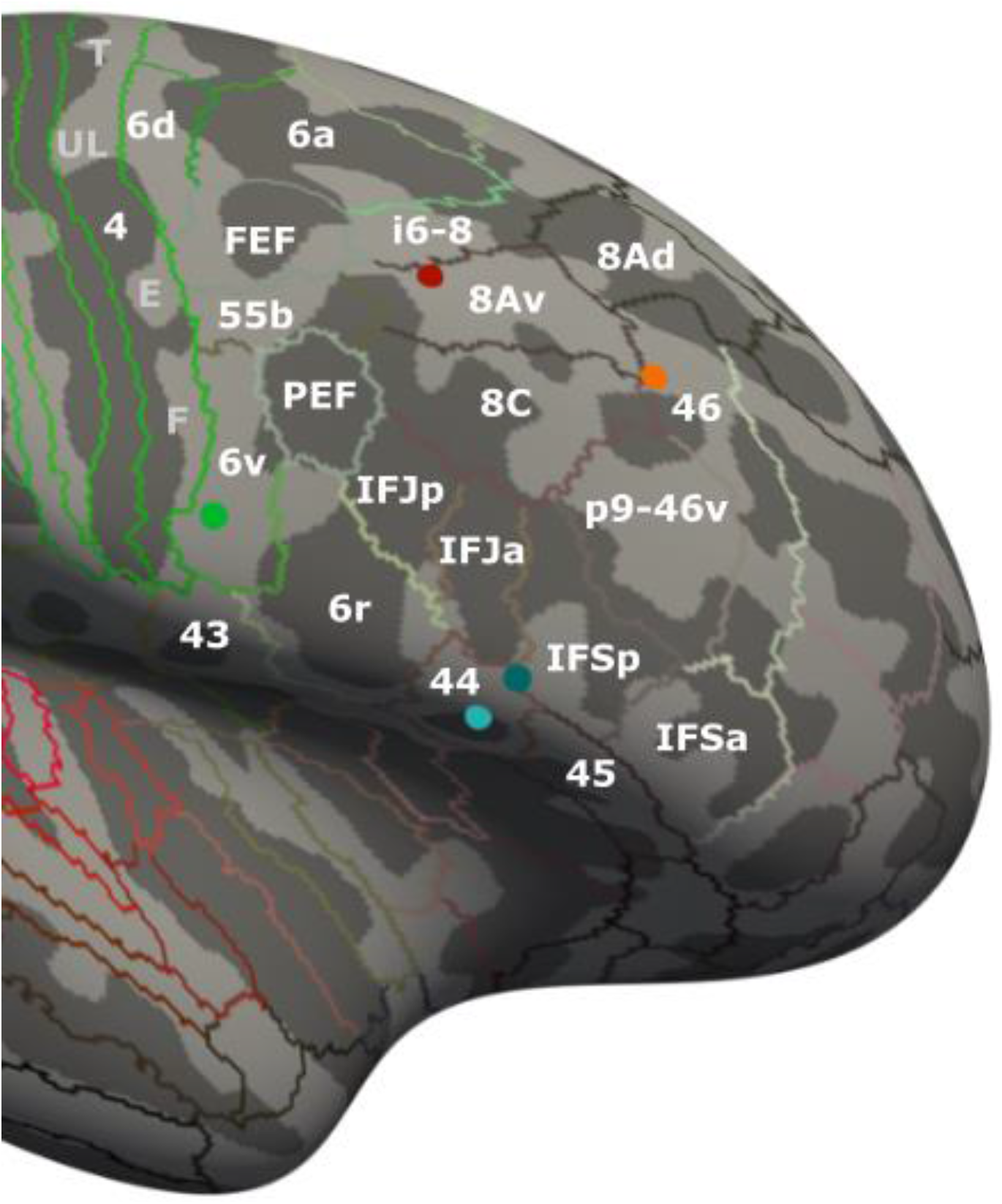
One-way ANOVA cluster centroids. The Human Connectome Project brain atlas (Glasser et al., 2016) was mapped onto the Freesurfer average brain to identify subdivisions of the frontal cortex. The significant cluster centroids were plotted on the Freesurfer brain using MNI coordinates (Table 1); red - area 8Av, orange - area 46, green - area 6v, light blue – 44, and cyan – area 44.

**Figure S2.**
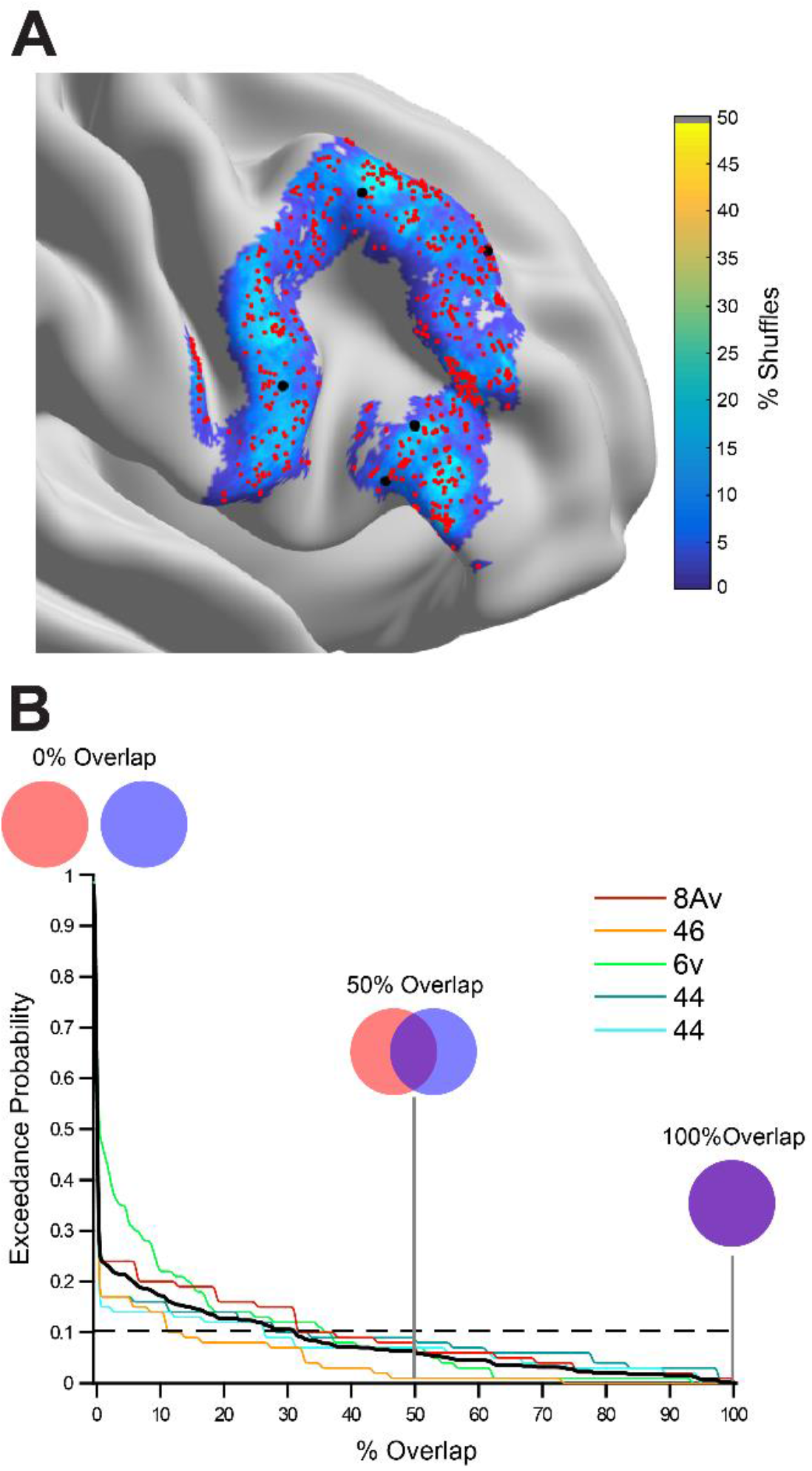
Shuffle analysis. **A.** For each vertex, the percent of shuffles in which it was identified as part of a cluster. Centroids of a clusters identified by the shuffled analysis are shown as red dots. The black dots show the centroids of the 5 clusters identified by the main analysis. **B.** The coloured lines show the exceedance probability for each of the original 5 clusters: the proportion of shuffles with clusters that overlap the original cluster by at least a given percentage. The average exceedance probability across clusters is shown as a thick black line. The horizontal dashed line shows the percentage of overlap at a threshold exceedance probability of 0.1.

